# Thioredoxin regulates the redox state and the activity of the human tRNA ligase complex

**DOI:** 10.1101/2023.05.26.542437

**Authors:** Dhaarsini Jaksch, Johanna Irnstorfer, Petra-Franziska Kalman, Javier Martinez

## Abstract

The mammalian tRNA ligase complex (tRNA-LC) catalyzes the splicing of intron-containing pre-tRNAs in the nucleus and the splicing of *XBP1* mRNA during the unfolded protein response (UPR) in the cytoplasm. We recently reported that the tRNA-LC co-evolved with PYROXD1, an essential oxidoreductase that protects the catalytic cysteine of RTCB, the catalytic subunit of the tRNA-LC, against aerobic oxidation. In this study we show that the oxidoreductase Thioredoxin (TRX) preserves the enzymatic activity of RTCB under otherwise inhibiting concentrations of oxidants. TRX physically interacts with oxidized RTCB, and reduces and re-activates RTCB through the action of its redox-active cysteine pair. We further show that TRX interacts with RTCB at late stages of UPR. Since the interaction requires oxidative conditions, our findings suggest that prolonged UPR generates reactive oxygen species. Thus, our results support a functional role for TRX in securing and repairing the active site of the tRNA-LC, thereby allowing pre-tRNA splicing and UPR to occur when cells encounter mild, but still inhibitory levels of reactive oxygen species.

## Introduction

RNA molecules are mostly synthesized as precursors and only become functional upon a series of modifications and processing events such as the removal of intronic sequences during a process called RNA splicing. The so-called “canonical splicing” entails the removal of introns from precursor mRNAs (pre-mRNAs) (Berget et al. 1977; Chow et al. 1977). In contrast, the term “non-canonical splicing” has been coined to describe the removal of single introns from pre-tRNAs (Hopper and Banks 1978; Knapp et al. 1978; O’Farrell et al. 1978; De Robertis and Olson 1979; Knapp et al. 1979; Peebles et al. 1979; Peebles et al. 1983; Schmidt and Matera 2020; reviewed in Phizicky and Hopper 2023).

Pre-tRNA splicing occurs in two steps: cleavage of the pre-tRNA by a tRNA splicing endonuclease (Ho et al. 1990; Rauhut et al. 1990; Trotta et al. 1997; Paushkin et al. 2004; Hayne et al. 2022; Sekulovski et al. 2022; Zhang et al. 2023) and ligation of the resulting tRNA exon halves by an RNA ligase to generate a mature tRNA (Filipowicz and Shatkin 1983; Greer et al. 1983; Laski et al. 1983; Phizicky et al. 1986). In mammals, the ligation of tRNA exon halves is executed by the tRNA ligase complex (tRNA-LC), composed of the catalytic subunit RTCB and four additional subunits: DDX1, CGI-99 (RTRAF/hCLE), FAM98B, and Ashwin (ASW/C2orf49) (Popow et al. 2011; Popow et al. 2012). The tRNA-LC becomes a multiple turnover enzyme through guanylylation by Archease (Popow et al. 2014; Desai et al. 2015). Splicing of intron-containing pre-tRNAs is essential because specific tRNA isodecoder families are exclusively encoded as intron-containing pre-tRNAs and therefore need to be spliced in order to be functional in protein translation (Chan and Lowe 2016; Gogakos et al. 2017).

Another example of non-canonical splicing is the cytoplasmic removal of a retained intron from an mRNA during the unfolded protein response (UPR), a signaling cascade that is triggered when unfolded proteins accumulate in the ER (Kozutsumi et al. 1988; Walter and Ron 2011). This pathway aims re-establish ER homeostasis to ensure survival of the cell, but can also induce apoptosis if activated over a prolonged period of time. The most conserved branch of the UPR comprises IRE1, an RNA endonuclease and kinase, that cleaves the retained intron in the *XBP1u* mRNA (u for unspliced) upon activation (Sidrauski and Walter 1997; Yoshida et al. 2001). In metazoans, the cleaved *XBP1* mRNA exons are joined by the tRNA-LC to generate *XBP1s* mRNA (s for spliced) (Jurkin et al. 2014; Kosmaczewski et al. 2014; Lu et al. 2014). Translation of this mRNA results in the transcription factor XBP1s, which upregulates proteins that assist in resolving the UPR (Lee et al. 2003; Acosta-Alvear et al. 2007). Besides these well-studied RNA ligation reactions, the tRNA-LC was found to be required for the splicing of transcripts related to DNA methylation and DNA damage repair in *Mus musculus* (Zhang et al. 2022).

We have recently reported that the mammalian tRNA-LC is reversibly inhibited by oxidative stress (Asanovic et al. 2021). Structural studies showed that the cysteine in the active site of an archaeal RTCB is prone to oxidation (Banerjee et al. 2021), suggesting that the inactivation of the mammalian tRNA-LC likely also occurs through oxidation of the catalytically active cysteine. Furthermore, we have shown that the oxidoreductase PYROXD1 co-evolved with the tRNA-LC to protect its catalytic center against aerobic oxidation, but not to rescue the tRNA-LC once oxidized (Asanovic et al. 2021). How the oxidized and inactivated tRNA-LC is re-activated remains unknown, as much as the physiological relevance of its regulation by a redox mechanism. Reduction of oxidized proteins during recovery from oxidative stress in cells is usually achieved by oxidoreductases such as Thioredoxin (TRX) (Luthman and Holmgren 1982; Lee et al. 2014). TRX reduces its target proteins using two catalytic cysteines – C32 and C35 in humans – within the CxxC motif. A proteome-wide study aiming to find targets of mammalian TRX and TRX-like proteins identified RTCB amongst other targets (Nakao et al. 2015). However, it remains unclear if this interaction has any physiological relevance.

In this study, we show that the absence of TRX enhances the sensitivity of the tRNA-LC towards oxidative stress, with an impact on pre-tRNA and *XBP1* mRNA splicing. We demonstrate that TRX re-activates the tRNA-LC after alleviation of the oxidative stress as well as upon extended UPR through direct interaction with RTCB.

## Results and Discussion

### TRX increases the tolerance of the tRNA ligase complex to oxidative stress

To test whether TRX preserves the enzymatic activity of the human tRNA-LC during oxidative stress, we generated a HeLa cell line expressing a Doxycycline (Dox) inducible short hairpin RNA (shRNA) targeting the *TXN* mRNA encoding TRX (shTRX cells) (Zuber et al. 2011). Both TRX protein and *TXN* mRNA levels decreased after 3 days of shRNA expression, while RTCB protein levels remained unchanged **(Suppl. Fig. 1A, B).** We monitored tRNA-LC activity through the inter-strand ligation of a double-stranded, radio-labeled RNA substrate bearing a 3’phosphate (3’P) on one strand and a 5’hydroxyl group (5’OH) on the complementary strand **(Fig. 1A)** (Popow et al. 2011). Under standard conditions, extracts from shTRX cells and control shRNA cells (shCtrl cells) displayed similar ligation activity **(Fig. 1A, B).** However, upon treatment with increasing concentrations of the oxidant menadione, the tRNA-LC became inactive at lower concentrations of menadione in shTRX cells **(Fig. 1C, D)**. We calculated IC_50_ values of 31,59 µM in shCtrl cells and 19,86 µM in shTRX cells. Treating cells with increasing concentrations of H_2_O_2_ caused a similar effect **(Suppl. Fig. 1C, D)**, with IC_50_ values of 75,84 µM for shCtrl cells and 37,21 µM for shTRX cells. RTCB levels remained unchanged in both treatments **(Suppl. Fig. 1E, F).** Of note, we observed an increase in the activity of the tRNA-LC irrespective of the presence of TRX at 10 µM menadione, but not in the presence of H_2_O_2_. This effect may relate to other conserved cysteine residues in the subunits of the tRNA-LC which are potentially modified during oxidative stress. Yet, the absence of TRX has a clear impact on the sensitivity of the tRNA-LC in the presence of menadione and H_2_O_2_.

**Figure 1:**
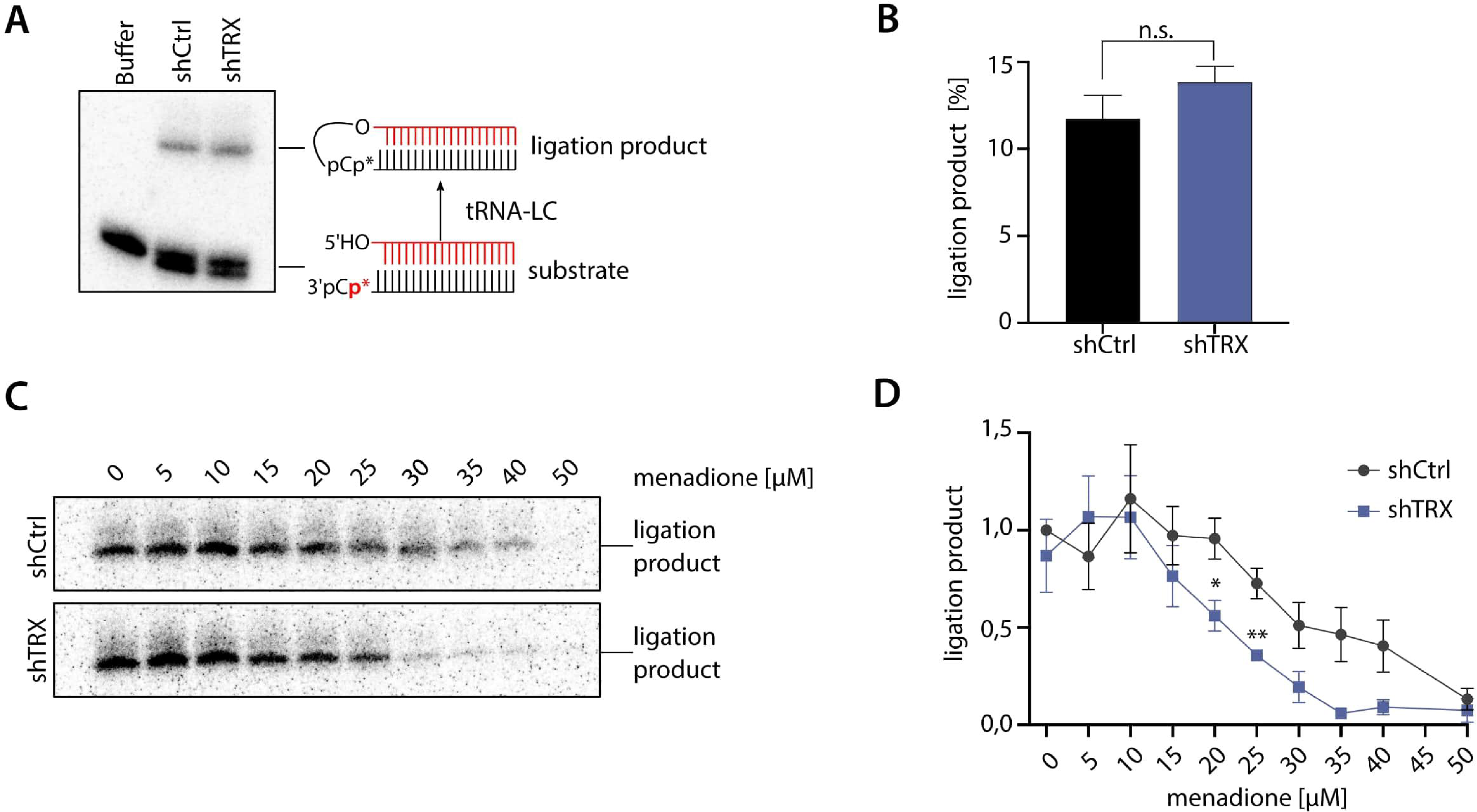
Depletion of TRX by shRNAs sensitizes the tRNA-LC to oxidative stress. A) HeLa cells expressing a short RNA hairpin (shRNA) targeting *TXN* mRNA to deplete TRX protein (shTRX) or a control shRNA targeting Renilla luciferase (shCtrl) upon addition of 2 µg/mL Doxycycline (Dox) were lysed after 3 days of shRNA expression and RNA ligation activity was assayed by incubating cell lysates with a double-stranded, radio-labeled RNA substrate containing a 3’ phosphate on one strand and a 5’OH on the complementary strand. Products of the reaction were resolved by denaturing urea polyacrylamide gel electrophoresis and visualized by phosphorimaging. B) Quantification of band intensities from Fig. 1A. The substrate conversion rate was calculated as the quotient of the substrate and the sum of substrate and product band intensities. (n=7, mean values ± SEM are shown, significances were analyzed using unpaired Student’s t test assuming unequal variances, with ∗, p < 0.05; ∗∗, p < 0.01; ∗∗∗, p < 0.005). C) ShCtrl and shTRX cells were treated with increasing concentrations of menadione for 1 h before harvest and lysis. Cell lysates were assayed as in Fig. 1A. D) Quantification of band intensities from Fig. 1C. The substrate conversion rate was calculated as in Fig. 1B, with n=3.

### Catalytically active TRX is required to sustain tRNA-LC activity during oxidative stress and to re-activate the tRNA-LC once the stress is mitigated

We next evaluated whether the catalytic cysteine residues of TRX are required to support tRNA-LC activity during oxidative stress. We therefore overexpressed shRNA-resistant, FLAG-tagged variants of human TRX – FLAG-TRX CxxC (WT) and FLAG-TRX SxxS (mutant) – in shTRX cells **(Suppl. Fig. 2A, B).** Overexpression of FLAG-TRX CxxC **(Suppl. Fig. 2B)** prevented the inactivation of the tRNA-LC by menadione **(Fig. 2A, B)**, while overexpression of FLAG-TRX SxxS did not **(Suppl. Fig. 2B, Fig. 2A, B)**. This shows that the catalytic cysteines of TRX are required to sustain the enzymatic activity of the tRNA-LC during oxidative stress.

**Figure 2:**
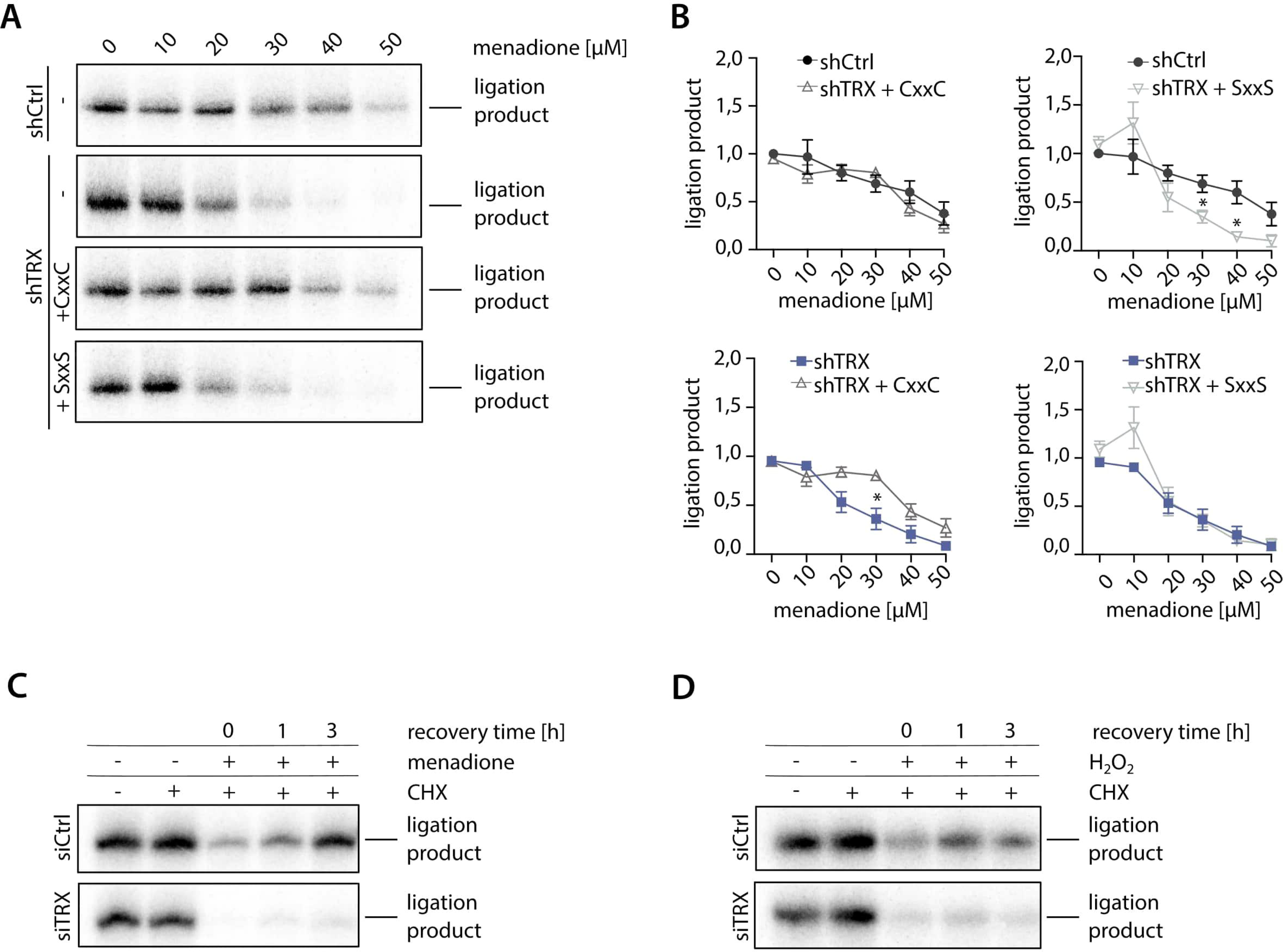
Catalytically active TRX is required to maintain tRNA-LC activity during oxidative stress and for its re-activation after oxidative stress levels decrease. A) ShRNA-resistant wildtype FLAG-TRX CxxC (labeled as CxxC) or active site double mutant FLAG-TRX SxxS (labeled as SxxS) were constitutively overexpressed in shTRX cells. Silencing of endogenous TRX was induced by addition of 2 µg/mL Dox for 3 days. Cells were treated with menadione for 1 h before harvest and lysis and RNA ligation activity was assayed by incubating cell lysates with a double-stranded, radio-labeled RNA substrate containing a 3’ phosphate on one strand and a 5’OH on the complementary strand. Products of the reaction were resolved by denaturing urea polyacrylamide gel electrophoresis and visualized by phosphorimaging. B) Quantification of band intensities from 2A. The substrate conversion rate was calculated as the quotient of the substrate and the sum of substrate and product band intensities and normalized to untreated shCtrl sample (n=3, mean values ± SEM are shown. Significances were analyzed using unpaired Student’s t test assuming unequal variances, with ∗, p < 0.05; ∗∗, p < 0.01; ∗∗∗, p < 0.005. Top left graph: comparison of ligation product in shCtrl vs. shTRX FLAG-TRX CxxC cells. Top right graph: comparison of ligation product in shCtrl vs. shTRX FLAG-TRX SxxS cells. Bottom left graph: comparison of ligation product in shTRX vs. shTRX FLAG-TRX CxxC cells. Bottom right graph: comparison of ligation product in shTRX vs. shTRX FLAG-TRX SxxS cells). C) HeLa cells were transfected with siRNA pools targeting *TXN* mRNA (siTRX) or a control siRNA (siCtrl) for 3 days. Cells were treated with 30 µM menadione for 30 min, washed, and allowed to recover for 0, 1 or 3 h in medium containing 10 µg/mL CHX. As a control, cells were either left untreated or treated with CHX alone for 5 h. Cells were lysed and assayed for RNA ligation as in Fig. 2A. D) HeLa cells were transfected with siRNA pools targeting *TXN* mRNA (siTRX) or a control siRNA (siCtrl) for 3 days. Cells were treated with 125 µM H_2_O_2_ for 30 min, washed, and allowed to recover for 0, 1 or 3 h in medium containing 10 µg/mL CHX. As a control, cells were either left untreated or treated with CHX alone for 5 h. Cells were lysed and assayed for RNA ligation as in Fig. 2A.

We next investigated the potential role of TRX in the previously described reversibility of the tRNA-LC inhibition upon recovery from oxidative stress (Asanovic et al. 2021), and in the re-activation the inhibited tRNA-LC. These experiments were performed in HeLa cells transiently transfected with siRNAs, instead of the Dox-inducible shRNA system, to avoid the antioxidant effect displayed by Dox (Clemens et al. 2018), which can affect the re-activation phase. We first performed titrations to determine the minimal inhibitory concentrations of menadione and H_2_O_2_ in siCtrl and siTRX cells. We observed an almost complete inhibition of tRNA-LC activity at 30 µM menadione in siCtrl cells, while tRNA-LC activity was abrogated at 20 µM menadione in siTRX cells **(Suppl. Fig. 2C)**. Inhibition with H_2_O_2_ was achieved at 250 µM in siCtrl cells, while 100 µM substantially reduced RNA ligation activity in siTRX cells **(Suppl. Fig. 2D)**.

To monitor the re-activation of the tRNA-LC after oxidative stress, we treated cells with 30 µM menadione, while further titrations with H_2_O_2_ led us to an optimal concentration of 125 µM H_2_O_2_. The treatment was done for 30 min and cells were allowed to recover over a time course of 3 h in medium supplemented with cycloheximide (CHX) to inhibit *de novo* protein synthesis. In siCtrl cells treated with menadione, the tRNA-LC fully recovered after 3 h **(Fig. 2C)**. In contrast, we did not detect tRNA-LC activity in siTRX cells **(Fig. 2C)**. Similar to menadione treatment, we were not able to detect tRNA-LC activity in siTRX cells treated with 125 µM H_2_O_2_ **(Fig. 2D)**. Importantly, levels of RTCB did not change during the recovery period after menadione or H_2_O_2_ treatment **(Suppl. Fig. 2E, F)**. These results indicate that catalytically active TRX is required to re-activate the tRNA-LC once oxidative stress levels have decreased.

### RTCB, the catalytic subunit of the tRNA-LC, is a substrate of TRX

We performed kinetic trapping pulldowns to test whether the reductive function of TRX relies on a physical interaction with the tRNA-LC (Nakao et al. 2015). We overexpressed FLAG-TRX CxxS in cells depleted of endogenous TRX to stabilize a potential interaction between the tRNA-LC and FLAG-TRX CxxS. Through mutation of the resolving C35, targets of TRX that have an oxidized cysteine residue (e.g. −SOH or a disulfide bond between two cysteines) are “trapped” due to the formation of an intermediate disulfide bond that cannot be reduced **(Fig. 3A).** We immunoprecipitated FLAG-TRX CxxS from untreated cells and from cells treated with different concentrations of H_2_O_2_ for 3 min, and carried out elution with FLAG-peptide in non-reducing conditions. We found RTCB, but no other subunits of the tRNA-LC, enriched in the eluate in a H_2_O_2_ concentration-dependent manner, indicating that RTCB is a target of TRX **(Fig. 3B).** Peroxiredoxin 1 (PRDX1), a well-known target of TRX (Chae et al. 1999), was also detected in the eluate **(Fig. 3B)**. A similar enrichment for RTCB and PRDX1 was observed in cells treated with increasing concentrations of menadione **(Fig. 3C)**.

**Figure 3:**
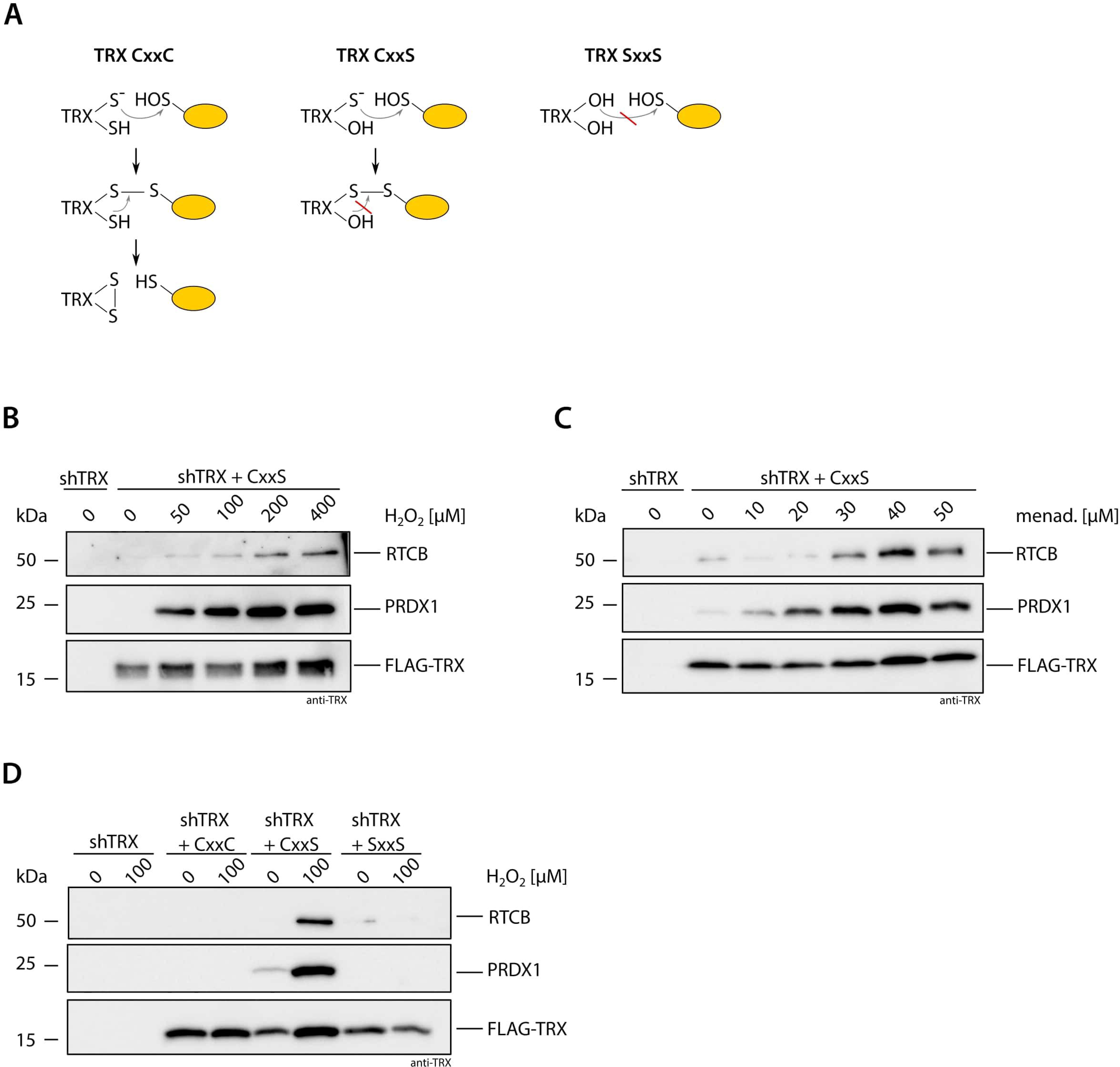
RTCB is a substrate of TRX. A) Scheme of the reduction mechanism of TRX CxxC towards its target protein containing an oxidized cysteine residue (left panel). Trapping of target proteins using TRX CxxS (middle panel). The double mutant SxxS cannot reduce the oxidized cysteine on the target protein due to mutation of the catalytic cysteines (right panel). B) shTRX cells constitutively overexpressing FLAG-TRX CxxS were treated for 3 min with increasing concentrations of H_2_O_2_ before lysis. ShTRX cells were left untreated as a negative control. Immunoprecipitation and elution were performed in non-reducing conditions. Eluates were analyzed by western blotting using anti-RTCB, anti-PRDX1 and anti-TRX antibodies. C) Trapping was performed as described in Fig. 3B in cells expressing shTRX FLAG-TRX CxxS after treatment of cells with indicated concentrations of menadione for 15 min. ShTRX were left untreated as a negative control. D) Trapping was performed as described in Fig. 3B in shTRX cells expressing FLAG-TRX CxxC, FLAG-TRX CxxS and FLAG-TRX SxxS after treatment of cells with 0 or 100 µM H_2_O_2_ for 3 min.

We next performed the trapping with FLAG-TRX CxxC in untreated and treated cells, and could not detect interaction after treatment with H_2_O_2_ due to immediate reduction of the intermediate disulfide bond by C35 **(Fig. 3D)**. When using the FLAG-TRX SxxS mutant, no trapping was observed, which is in line with the fact that C32 is required for formation of a mixed disulfide bond **(Fig. 3D).**

Thus, our data demonstrate that TRX directly interacts with RTCB in an oxidant-dependent manner. This suggests that the increased redox sensitivity of the tRNA-LC in the absence of TRX, as well as the reversible character of the inactivation, is a consequence of the interaction between TRX and RTCB. Furthermore, these results strongly imply that the re-activation of RTCB is through the reductive function of TRX, presumably by reduction of the active site cysteine of RTCB. This is in line with the structural study from *Pyrococcus horikoshii*, where oxidation of the active site cysteine was observed in aerobically purified RTCB (Banerjee et al. 2021). Oxidation of the highly conserved cysteine (C122 in humans) causes loss of metal ion binding and therefore loss of catalytic activity. We hypothesize that TRX can reduce C122 and thereby allows re-loading of the active site with the metal ions to regain its function.

### Depletion of TRX during oxidative stress impairs pre-tRNA splicing and XBP1 mRNA splicing during UPR

We tested the TRX-RTCB interplay during pre-tRNA splicing and *XBP1* mRNA splicing in conditions of oxidative stress. For this purpose, we assayed pre-tRNA splicing *in vitro* using a radio-labeled, intron-containing pre-tRNA from *S. cerevisiae* and lysates from shCtrl and shTRX cells treated with menadione. Cleavage of the substrate pre-tRNA with the consequent generation of tRNA exon halves was not impaired by oxidative stress **(Fig. 4A, bottom panels)** (Asanovic et al. 2021). However, ligation of exon halves to generate a mature tRNA was inhibited at 25 µM menadione in shTRX cells, compared to 50 µM menadione in shCtrl cells **(Fig. 4A, top panels)**.

**Figure 4:**
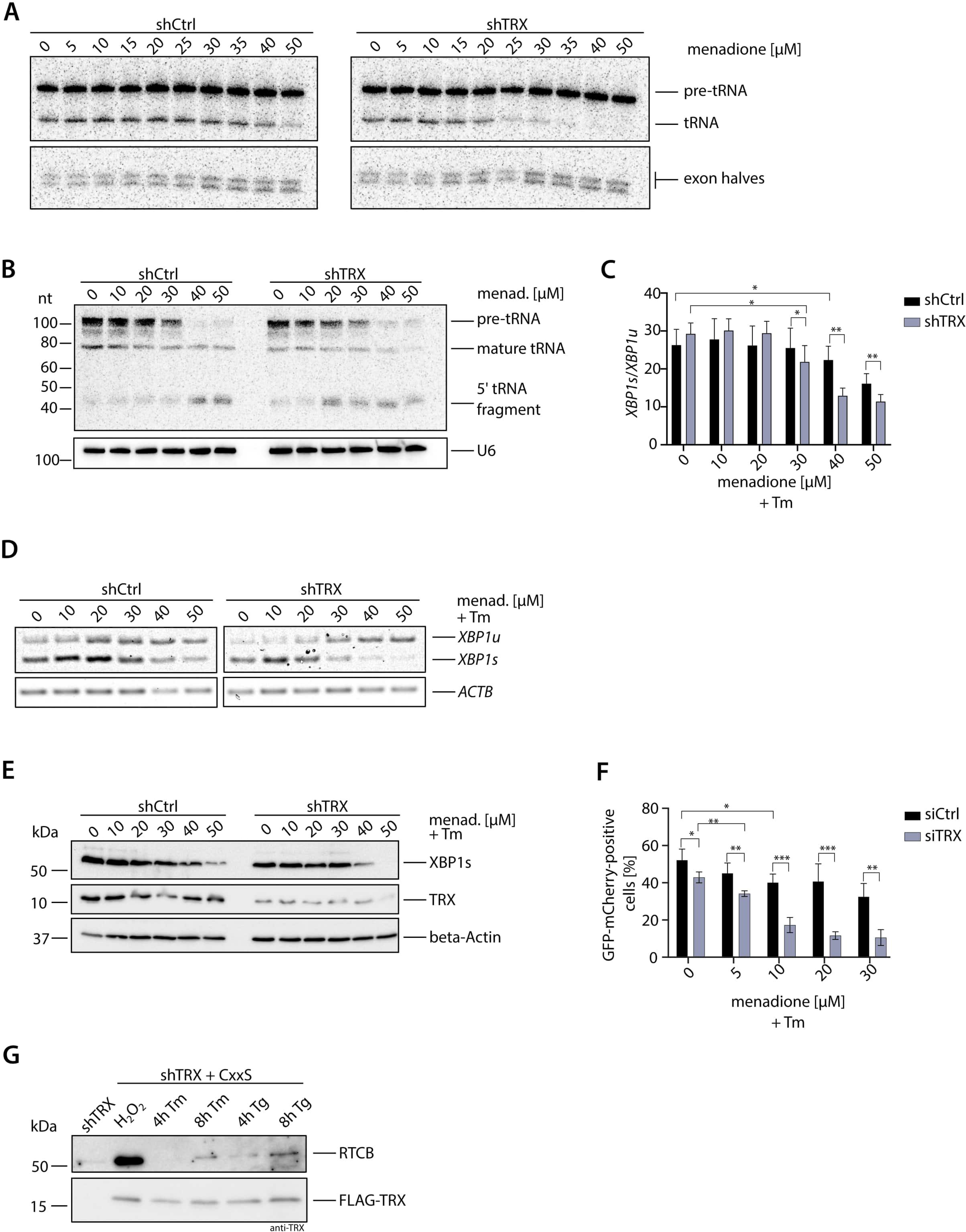
Depletion of TRX impairs physiological functions of the tRNA-LC. A) ShCtrl and shTRX cells were treated with indicated menadione concentrations for 1 h before harvest and lysis. An internally radio-labeled *S. cerevisiae* pre-tRNA substrate was added to cell lysates to monitor cleavage by the TSEN complex and ligation to mature tRNAs by the tRNA-LC. Pre-tRNA processing was monitored by denaturing urea gel electrophoresis and visualized by phosphorimaging. B) ShCtrl and shTRX were treated with menadione for 1 h. RNA was isolated and subjected to northern blot analysis using a probe complementary to the 5’exon of Tyr-tRNA. A probe targeting U6 snRNA was used as a loading control. C) ShCtrl and shTRX cells were treated with indicated menadione concentrations and 1,5 µg/ml Tm for 4 h before RNA isolation and cDNA synthesis. RT-qPCR was performed to measure *XBP1s* and *XBP1u* mRNA levels relative to *ACTB* (shown as the ratio *XBP1s/XBP1u*). (n=9, mean values ± SEM are shown. Significances were analyzed using paired Student’s t test assuming unequal variances, with ∗, p < 0.05; ∗∗, p < 0.01; ∗∗∗, p < 0.005). D) ShCtrl and shTRX cells were treated with indicated menadione concentrations and 1,5 µg/ml Tm for 4 h before RNA isolation and cDNA synthesis. Semi-quantitative RT-PCR was performed to analyze levels of *XBP1* mRNA and *ACTB* mRNA. PCR products were resolved by agarose gel electrophoresis. E) ShCtrl and shTRX cells were treated with indicated menadione concentrations and 1,5 µg/mL Tm for 4 h before harvest. Cells were lysed and protein levels of XBP1s and TRX were analyzed by western blotting using respective antibodies. Beta-Actin levels were assessed as a loading control. F) HEK293 FITR cells expressing GFP-XBP1^intron^-mCherry were transfected with siRNA pools targeting *TXN* (siTRX) or control siRNA (siCtrl) for 3 days. Expression of the reporter was induced for 5 h with 2 µg/ml Dox and treatment with menadione and 1,5 µg/mL Tm was performed for 4 h. Splicing of the intron (generation of GFP-mCherry fusion protein) was assessed by FACS analysis. The graph only shows GFP-mCherry positive cells (n=4, mean values ± SEM are shown. Significances were analyzed using unpaired Student’s t test assuming unequal variances, with ∗, p < 0.05; ∗∗, p < 0.01; ∗∗∗, p < 0.005). G) ShTRX FLAG-TRX CxxS cells were treated for 4 or 8 h with 1,5 µg/mL Tm, or for 4 or 8 h with 300 nM Tg before lysis. As a positive control for trapping, shTRX FLAG-TRX CxxS cells were treated for 3 min with 100 µM H_2_O_2_. ShTRX cells were left untreated as a negative control. Immunoprecipitation and elution were performed in non-reducing conditions. Eluates were analyzed by western blotting using anti-RTCB or anti-TRX antibodies.

To further validate the protective role of TRX towards the tRNA-LC under oxidative stress conditions, we performed a northern blot analysis to detect a fragment of Tyr-tRNA, composed of the 5’ leader sequence followed by the 5’ exon, that accumulates upon inhibition of the tRNA-LC (Hanada et al. 2013). The Tyr-tRNA fragment was detected at lower concentrations of menadione in shTRX cells compared to shCtrl cells **(Fig. 4B)**. Of note, levels of total pre-tRNAs after treatment with 40 and 50 µM menadione were reduced in both cell lines, possibly due to Pol III inhibition (Gouge et al. 2015).

We next monitored *XBP1* mRNA splicing upon induction of UPR with Tunicamycin (Tm) and simultaneous oxidative stress conditions. The splicing efficiency, determined by the ratio of *XBP1s* to *XBP1u* mRNA, was decreased in shCtrl and shTRX cells after menadione treatment when measured by RT-qPCR **(Fig. 4C)**. However, in shTRX cells, 30 µM menadione was sufficient for a significant reduction in comparison to shCtrl cells, where treatment with 40 µM menadione was required. A similar effect was observed when measuring levels of *XBP1u* and *XBP1s* mRNA by semi-quantitative RT-PCR **(Fig. 4D)**. We also analyzed levels of XBP1s protein by western blot and observed a reduction at 40 µM menadione in shTRX cells, while 50 µM menadione was required in shCtrl cells to achieve a similar effect **(Fig. 4E)**. The reduction in *XBP1s* mRNA and XBP1 protein levels in the absence of TRX and simultaneous oxidative stress also affected XBP1s target genes, as shown by the reduction of *DNAJB9* mRNA in shTRX cells treated with 30 µM menadione in comparison with shCtrl cells **(Suppl. Fig. 3A)**.

We developed a FACS-based *XBP1* mRNA splicing reporter in a HEK293 FITR cell line. The reporter encodes a Dox-inducible, GFP-XBP1^intron^-mCherry transcript that generates a GFP-mCherry fusion protein upon splicing, similar to previously described XBP1 splicing reporters (Lu et al. 2014; Roy et al. 2017). First, we validated the functionality of the reporter by co-depleting RTCB and Archease, which was shown to be required for a decrease in *XBP1s* generation (Jurkin et al. 2014). We observed a drastic reduction in the generation of GFP-mCherry when UPR was induced with Thapsigargin (Tg) in cells depleted of RTCB and Archease, demonstrating that the reporter indeed reflects endogenous *XBP1* mRNA splicing **(Suppl. Fig. 3B)**. When TRX was depleted using siRNAs (siTRX cells) and UPR was induced simultaneously with oxidative stress, the levels of GFP-mCherry protein diminished at lower menadione concentrations in comparison to siCtrl cells **(Fig. 4F)**. The difference in the concentrations of menadione required to observe a decrease in the expression of the GFP-mCherry reporter and the endogenous XBP1 protein in shCtrl/shTRX cells is due to the use of different cell lines: HEK293 FITR for the reporter and HeLa cells for the shRNA-mediated silencing. Taken together, these results demonstrate that TRX is required to maintain the physiological activity of the tRNA-LC when cells encounter oxidative stress.

We finally took advantage of the sensitive trapping approach to test whether the physical interaction between TRX and the tRNA-LC also occurs as a consequence of ROS generated endogenously during UPR. We therefore treated cells for 4 or 8 h with Tm or Tg to induce UPR. As a positive control for the trapping, we treated cells with 100 µM H_2_O_2_ **(Fig. 3B)**. We detected a robust interaction between FLAG-TRX CxxS and RTCB when treating cells with Tg for 8 h **(Fig. 4G)**. This result suggests that ROS are generated during prolonged UPR and that RTCB is supported by the reductive activity of TRX. The tRNA-LC catalyzes the splicing of *XBP1* mRNA at low concentrations of exogenous ROS in the presence of TRX **(Fig. 4C-F);** however, TRX cannot cope with higher levels of oxidation, leading to a decrease in *XBP1* mRNA splicing and, consequently, in the expression of XBP1s target genes **(Suppl. Fig. 3A).** Thus, potentially higher levels of ROS generated at late stages of UPR could irreversibly inhibit the tRNA-LC, downregulate *XBP1* mRNA splicing and signal to induce apoptosis.

This study demonstrates that the redox sensitivity of the tRNA-LC is a tightly regulated process that involves several factors and impacts pre-tRNA splicing, UPR and ultimately, protein biosynthesis. PYROXD1 has been shown to protect RTCB from oxidation (Asanovic et al. 2021) and, recently, to dissociates from RTCB before substrate binding (Loeff et al. 2023). Such dissociation offers an opportunity for ROS to oxidize RTCB. We envision that TRX acts at this step, bringing RTCB back to a functional state.

## Materials and Methods

### Cell culture

HeLa Kyoto, HeLa RIEP (Jurkin et al. 2014) and HEK293 Flip-In™ T-REx™ (FITR, Invitrogen) cells were cultured in Dulbecco’s modified Eagle’s medium (DMEM high glucose, Gibco, #41966-052) supplemented with 10% fetal bovine serum (FBS, Gibco, #A5256701), 100 µg/mL penicillin/100 µg/mL streptomycin (Sigma Aldrich, #P0781-100ml) and 20 mM HEPES pH 7,0 (Gibco, #15630-122), at 37°C, 5% CO_2_. Cells were regularly tested for mycoplasma contamination.

### Generation of Dox-inducible shRNA cells lines and induction of shRNA expression

For generation of Doxycycline-inducible shRNA cell lines, HeLa RIEP cells were transduced with ecotropically packaged shRNA expression vectors (pSIN-TRE3G-GFP-miRE-PGK-Neo, see ‘cloning of shRNAs’) or rescue vectors (pBMN-FLAG-IRES-GFP, see ‘cloning of shRNA-resistant constructs’) as described before (Fellmann et al. 2013). For retroviral packaging, Platinum E cells (PlatE, Cell Biolabs) were seeded into 10 cm^2^ plates and transfected at 90% confluency with 20 µg plasmid DNA and 10 µg helper plasmid (pCMV-Gag-Pol, Cell Biolabs) using CalPhos Mammalian Transfection Kit (Clontech®/Takara, #631312) according to manufacturer’s instructions. Transfection medium was replaced to complete DMEM after 16 h. Virus-containing supernatants were harvested at 36, 48 and 60 h post transfection and added dropwise onto HeLa RIEP cells for transduction. Cells expressing the shRNA constructs were selected using 1 mg/mL Neomycin (Calbiochem, #480100). To induce expression of shRNAs cells were treated for 3 days with 2 µg/mL Dox.

### *XBP1* splicing reporter

The *XBP1* splicing reporter was designed similar to previously described studies (Lu et al. 2014). An 120-mer oligonucleotide containing the *XBP1* intron was synthesized (Sigma Aldrich, sequence, intron in bold: 5’AGCCAAGGGGAATGAAGTGAGGCCAGTGGCCGGGTCTGCTGAGTCCGCAGCA**CTCAGACTACGT GCACCTCTGCAGCA**GGTGCAGGCCCAGTTGTCACCCCTCCAGAACATCTCCCCATG-3’), flanked by restriction sites and cloned into a pcDNA5-FRT vector containing EGFP and mCherry sequences. For generation of stable HEK293 FITR cells expressing the GFP-*XBP1*^intron^-mCherry reporter, cells were seeded into a 6-well plate and transfected at 50% confluency. 1,5 µg of plasmid DNA and 1,5 µg pOG44 (encoding the Flp recombinase) were mixed and transfected using CalPhos Mammalian Transfection Kit (Clontech®/Takara, #631312). The transfection medium was replaced by complete DMEM after 24 h, and after further 24 h, selection of positive clones was started using 0,5 mg/mL Hygromycin B. Colonies appeared after 14 days of selection. For UPR-related experiments, silencing of TRX or RTCB and Archease was done as described below. Expression of the GFP-XBP1^intron^-mCherry was induced for 5 h, 1 h later menadione and Tm were added to the cells. Cells were harvested by trypsinization, pellets were resuspended in FACS buffer (PBS + 2% FCS) and analyzed by flow cytometry (BD LSRFortessa™) and FlowJo software.

### Silencing of TRX using siRNA

For depletion of proteins using RNAi, cells were seeded one day before transfection. Small interfering RNAs (siRNA) targeting human *TXN* (siTRX) or scrambled control (siCtrl) consisted of siRNAs pools of 30 siRNAs and were purchased from siTOOL Biotech (Munich, Germany). HeLa cells were transfected at 30-40% confluency using Lipofectamine RNAiMax (Invitrogen, #13778-150) according to manufacturer’s instructions. Subsequent experiments were performed after 3 days of silencing.

### Treatment with inhibitors and UPR induction

For treatment with inhibitors or oxidants, HeLa Kyoto cells were seeded into 6-well plates and treated at 90% confluency. A list of compounds and inhibitors used in this work can be found in **Error! Reference source not found.**. For induction of the UPR in HeLa RIEP shRNA cells, knockdown of TRX was induced using 2 µg/mL Dox for 3 days. Cells were treated at 80% confluency with 300 nM Thapsigargin (Sigma Aldrich) or 1,5 µg/mL Tunicamycin (Sigma Aldrich).

### Cloning of shRNAs

For RNAi-mediated depletion of TRX, shRNAs targeting *TXN* mRNA were designed using the online siRNA prediction tool ‘Splash RNA’ (http://splashrna.mskcc.org) (Fellmann et al. 2013; Pelossof et al. 2017). The sequence of shCtrl and shTRX shRNA guides can be found in table 2. The respective 97-mer oligonucleotides (Sigma Aldrich, for guide sequences see table 2) were amplified by PCR using the following conditions: 0,5 ng of template DNA, 0,5 µL of Phusion polymerase (NEB, #M0530S), 1x buffer GC, 3% DMSO, 0,3 µM of each primer (miR30_fwd: 5’-CAG AAG GCT CGA GAA GGT ATA TTG CTG TTG ACA GTG AGC G-3’ and miR30_rev: 5’-CTA AAG TAG CCC CTT GAA TTC CGA GGC AGT AGG CA-3’). The PCR was performed using the following protocol: 95 °C for 2 min; 33 cycles of 95 °C for 15 s, 58 °C for 30 s and 72 °C for 25 s; 72 °C for 5 min. PCR products were purified using QIAquick® PCR Purification Kit (QIAGEN, #28104) according to manufacturer’s instructions. Purified PCR products and the destination vector pSIN-TRE3G-GFP-miRE-PGK-Neo were digested with XhoI/EcoRI 37°C for 1 h and the PCR product was ligated into the destination vector using T4 DNA ligase (NEB, # M0202L) overnight at 16 °C.

**Table 1:**
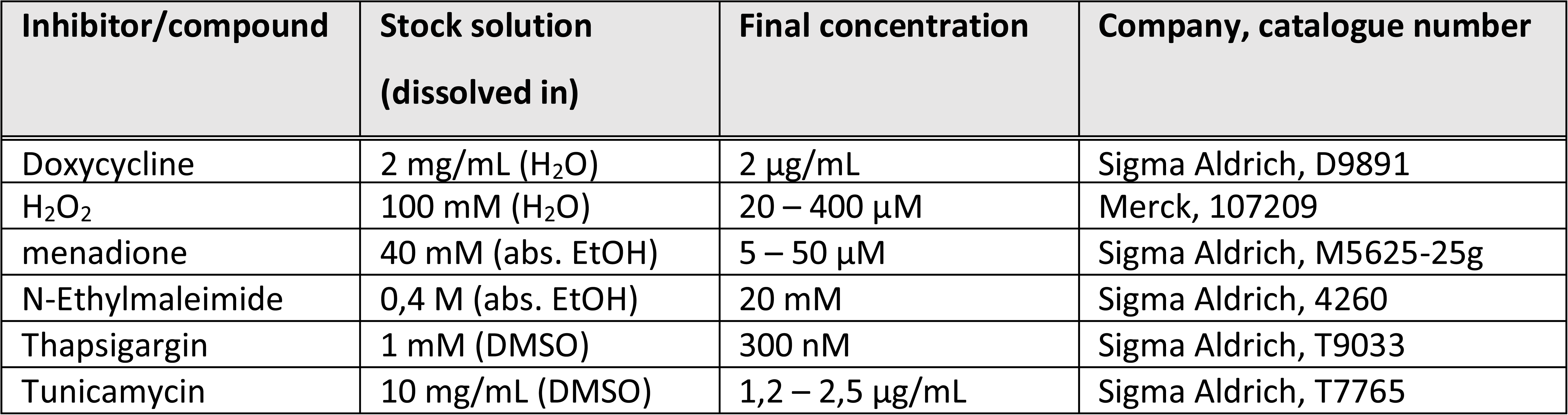
Inhibitors and compounds used in this study

**Table 2:**
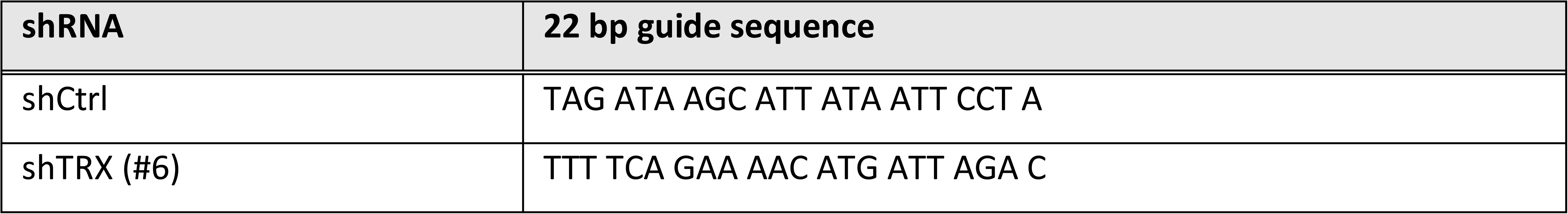
shRNA guide sequences

### Cloning of shRNA-resistant TRX-rescue constructs

The sequence of human TRX cDNA was adapted to be non-targetable by shTRX #6. The cDNA of WT (CxxC), C35S (CxxS) and the double mutant C32S/C35S (SxxS) was cloned into pDONR221 (Invitrogen) by Gateway recombination. Sequences were subsequently shuttled from the pDONR221 construct into pBMN-FLAG-IRES-GFP (introducing an amino-terminal FLAG epitope tag) that was used to generate stable cell lines. The primer sequences for Gateway recombination are shown in table 3 (small letters in primer sequence indicate the changed base for shRNA-resistance).

**Table 3:**
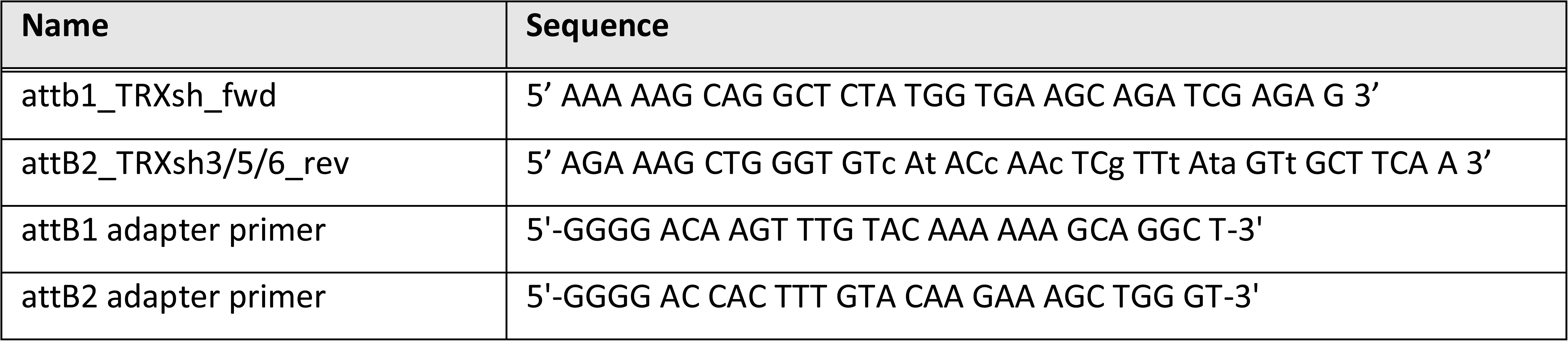
Primers used to generate shRNA-resistant TRX variants

### Transformation of *E.coli* and plasmid isolation

DNA constructs were mixed with 50 µL of chemically competent DH5α cells, and incubated at 4 °C for 30 min. The bacteria were heat shocked at 42 °C for 45 sec and incubated at 4 °C for 2 min. 450 µL of LB medium was added and incubated at 37°C, for 1 h while shaking. The bacteria were plated onto agar plates supplemented with the appropriate antibiotic and incubated overnight at 37 °C.

For plasmid isolation, one colony of transformed bacteria was inoculated into 5 mL (Mini prep) or 250 mL (Maxiprep) LB medium supplemented with the appropriate antibiotic and incubated overnight at 37 °C while shaking at 200 rpm. Isolation of plasmids was done using QIAprep® Miniprep or Maxiprep kits (Qiagen) according to manufacturer’s instructions. Purified plasmids were resuspended in nuclease-free water and DNA concentration and purity were measured at A_260nm_ using DeNovix DS-11 spectrophotometer.

### Preparation of whole cell extracts for activity assays

For tRNA-LC activity assays, cells were scraped with cold PBS and transferred into a tube. After spinning at 1000 rpm, 5 min, 4 °C, the supernatant was removed and the cell pellet resuspend in HeLa lysis buffer (10% glycerol, 30 mM HEPES pH 7,4, 5 mM MgCl_2_, 100 mM KCl, 1% Nonidet P-40, 0,1 mM AEBSF, 1 mM TCEP pH 7,0). Whole cell extracts were cleared by centrifugation at 14 8000 rpm, 10 min, 4 °C, the supernatant was transferred into a tube and protein concentration was measured by Bradford assay.

### Preparation of extracts for SDS-PAGE

For SDS-PAGE and western blot analysis, cells were scraped with cold PBS and transferred into a tube. After spinning at 1000 rpm, 5 min, 4 °C, the supernatant was removed, the cell pellet was resuspended in 5x SDS-loading buffer (62,5 mM Tris pH 6,8, 25 mM EDTA pH 8,0, 5% SDS, 5% β-mercaptoethanol, 0,025% bromophenol blue, 50% glycerol), diluted with water to obtain 1x SDS-loading buffer. Samples were boiled for 10 min at 95 °C and were frozen at −20 °C or subjected to protein analysis by SDS-PAGE and western blot.

### Determination of protein concentration by Bradford assay

For measurement of protein concentration of whole cell extracts, Bradford Protein Assay (Bio-Rad, #500-0006) was mixed 1: 5 in water. Concentration standards were prepared with bovine serum albumin (BSA, 1 mg/mL), by adding 0, 1, 2, 4, 8 or 16 µL of BSA to 1 mL of Bradford reagent and a standard curve was prepared by measuring the absorbance at 595 nm. 1 µL of cleared lysate was diluted in 1 mL Bradford reagent and concentrations were measured at 595 nm based on the standard curve using DeNovix DS-11 spectrophotometer.

### SDS-PAGE and western blot

25-30 μg of a sample was separated by SDS-PAGE and transferred onto methanol-activated Immuno-Blot® PVDF membranes in 1x Turbo Transblot buffer (Bio-Rad, #1704273). Blotting of two membranes was performed at 400 mA for 1 h. Membranes were blocked for 1 h with 5% milk in PBS with 0,05% Tween-20 (PBS-T) or 3% BSA in PBS-T. Blocked membranes were probed with primary antibodies diluted in 5% milk PBS-T or 3% BSA in PBS-T overnight at 4 °C while shaking. Membranes were washed three times in PBS-T for 5-10 min and incubated with the appropriate secondary antibody in 5% milk in PBS-T for 1 h at room temperature, before washing three times for 5-10 min in PBS-T. A list of primary and secondary antibodies and their dilutions used in this work can be found in table 4. Western blots were developed using Clarity™ or Clarity™ Max Western ECL substrate (Bio-Rad, # 170-5061 and #1705062) and visualized using ChemiDoc™ Gel Imaging System (Bio-Rad).

**Table 4:** Antibodies and dilutions used in this study

| Antibody | Dilution | Company, catalog number |
| --- | --- | --- |
| α-Actin, rabbit | 1:5 000 in 5% milk PBS-T | Sigma Aldrich, #A2066 |
| α-FLAG, mouse | 1:1 000 in 5% milk PBS-T | Sigma Aldrich, #F1804 |
| α-mouse HRP | 1:20 000 in 5% milk PBS-T | Invitrogen, #62-6520 |
| α-PRDX1, rabbit | 1:1 000 in 3% BSA PBS-T | Cell Signaling, #8499S |
| α-rabbit HRP | 1:20 000 in 5% milk PBS-T | Invitrogen, #65-6120 |
| α-RTCB, rabbit | 1:1 000 in 5% milk PBS-T | Bethyl, #A305-079A |
| α-TRX, rabbit | 1:1 000 in 3% BSA PBS-T | Cell Signaling, #2429 |
| α-XBP1s, rabbit | 1:1 000 in 3% BSA PBS-T | Abcam, #ab220783 |

### RNA isolation and cDNA synthesis

Isolation of RNA from cells was performed using TRIzol™ reagent (Invitrogen, #15596018) according to manufacturer’s instructions. The obtained RNA pellet was airdried and resuspended in 20 µL nuclease-free water. Concentration and purity of the sample was measured at A_260/280nm_ using DeNovix DS-11 spectrophotometer. Up to 2 µg RNA was treated with DNase to remove genomic DNA. cDNA synthesis from isolated RNA was prepared using Maxima First Strand cDNA Synthesis kit for RT-qPCR (Thermo Scientific, #K1672) according to manufacturer’s instructions. Obtained cDNA was diluted 1:10 in nuclease-free water before downstream analysis.

### Quantitative RT-PCR

For a quantitative measurement of mRNA levels using RT-qPCR, a master mix was prepared for each set of primers: for one well of a 384-well plate, 5 µL of 2x GoTaq (Promega, #A600A), 0,4 µL forward primer (0,5 µM), 0,4 µL reverse primer (0,5 µM) and 3,2 µL nuclease-free water were mixed. 1 µL of cDNA was pipetted into the 384-well plate, and 9 µL master mix was added per well. Primers used in this work can be found table 5. The RT-qPCR was performed in CFX384 Touch (Bio-Rad) using the following protocol: 50 °C for 10 min, 95 °C for 5 min, followed by 60 cycles in total at 95 °C for 10 sec and 60 °C for 30 sec. The quality of qPCR primers was evaluated by melting curve analysis and the obtained Ct values were analyzed using the ΔΔCt method.

**Table 5:**
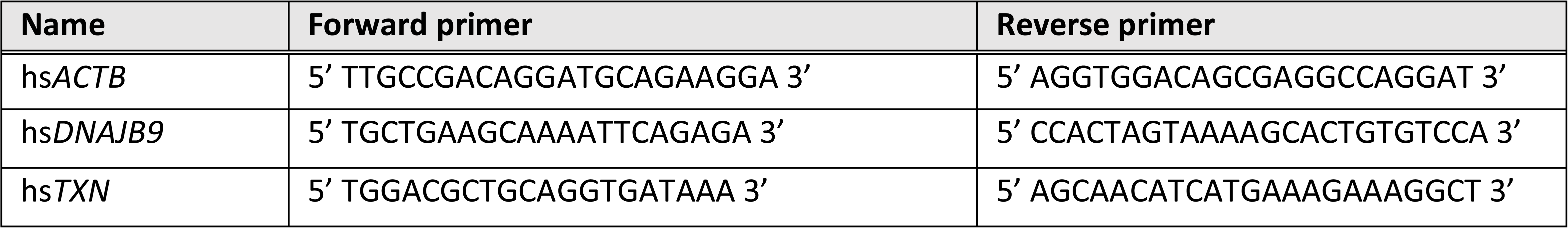
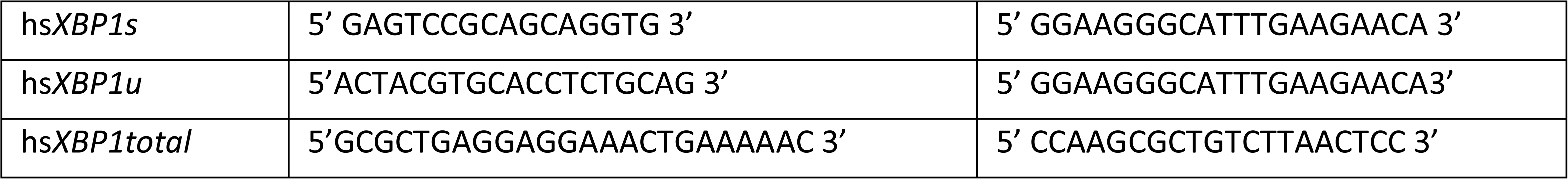
Primers used for RT-qPCR

### Semi-quantitative RT-PCR

For examination of *XBP1u* and *XBP1s* mRNA levels by semi-quantitative reverse transcription PCR (RT-PCR), 1 µL of diluted cDNA was mixed with 12,5 µL ReqTaq® Ready Mix™ PCR reaction mix (Sigma Aldrich, #R2523), 0,5 µL forward primer (0,2 µM), 0,5 µL reverse primer (0,2 µM) and filled up to 25 µL with nuclease-free water. The sequences for *XBP1* and *ACTB* forward and reverse primer pairs can be found in table 6 (Jurkin et al. 2014). DNA amplification was performed at 98 °C for 3 min, followed by 27 cycles of 98 °C for 30 s, 55 °C for 45 s and 72 °C for 1 min, and finalized by incubation at 72 °C for 10 min. The products of the PCR were run on a 3% agarose gel for 3 h at 50 V and quantified using ImageJ software.

**Table 6:**
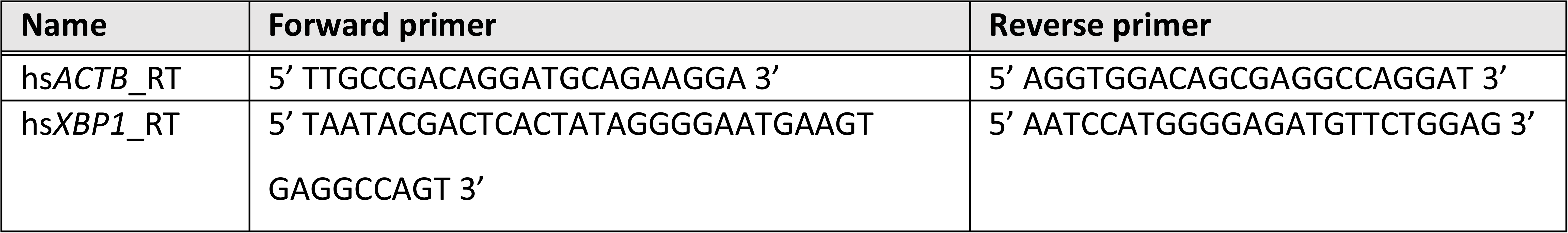
Primers used for semi-quantitative RT-PCR

### Northern blot analysis

For northern blot analysis of RNA samples, 5 µg of total RNA was mixed with equal volumes of 2x FA buffer (90% formamide, 50 mM EDTA, 1 ng/mL bromophenol blue, 1 ng/mL Xylene Cyanol) and boiled for 5 min at 98 °C. Samples were resolved using a 10% denaturing urea polyacrylamide gel (SequaGel®, National Diagnostics, #EC-833). After the gel run, the RNA was blotted onto Hybond® N+ membranes (Amersham, #RPN303B) for 3 h at 180 mA and cross-linked by UV. Pre-hybridization was carried out in hybridization buffer (5x SSC, 20 mM Na_2_HPO_4_ pH 7,2, 7% SDS, and 0,1 mg/mL sonicated salmon sperm DNA (Agilent Technologies, #201190)) at 50 °C for 1 h, and hybridization was done at 50 °C (DNA probe) or 80 °C (LNA probe) overnight in hybridization buffer supplemented with 100 pmol of [5’-^32^P] labeled DNA or LNA probes. To obtain the radioactive probe, 1 µL of the respective probe (100 µM) was labeled using 1 µL 10x PNK buffer, 2 µL [γ-^32^P]ATP, 1 µL PNK and 14 µL nuclease-free water. The reaction was incubated for 1 h at 37 °C. The labeled probe was separated from unincorporated nuclides using MicroSpin G-25 columns (GE healthcare, #GE27-5325-01) according to manufacturer’s instructions. The DNA/LNA probes used for northern blots can be found in table 7. Blots were washed twice for 1 min with 5x SSC, 5% SDS and once with 1x SSC, 1% SDS at 50 °C. The membrane was exposed and visualized by phosphorimaging. For re-probing, membranes were boiled for 5 min in 0,1% SDS and 0,1x SSC, pre-hybridized and probed with labeled probes as described before. Quantification of band intensities was performed using ImageQuant software (GE Healthcare).

**Table 7:**
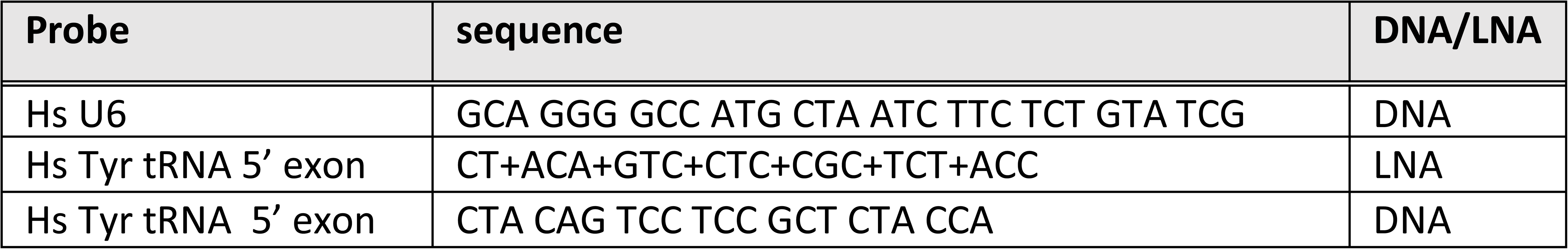
DNA or LNA probes used for northern blots

### Generation of radio-labeled inter-strand RNA substrate

1,11 MBq [5’-^32^P]cytidine-3’,5’-bisphosphate ([^32^P]Cp, 111 TBq/mmol, Hartmann Analytic) were ligated to 50 pmol RNA oligonucleotide “20.25” (5’-UCG AAG UAU UCC GCG UAC GU-3’, Dharmacon) with 1 µL T4 RNA ligase 1 (NEB, #M0204S) for 1 h at 16 °C in 15% (v/v) DMSO, 50 mM Tris-HCl pH 7,6, 10 mM MgCl_2_, 10 mM β-mercaptoethanol, 200 µM ATP, 0,1 mg/mL BSA in a total reaction volume of 20 µL. Labelling reactions were stopped with equal volumes of 2x FA buffer (90% formamide, 50 mM EDTA, 1 ng/mL bromophenol blue, 1 ng/mL Xylene Cyanol), boiled for 5 min at 98 °C and separated on 10% denaturing urea polyacrylamide gel (SequaGel®, National Diagnostics, #EC-833). Labeled RNA was visualized by autoradiography, cut from the gel and passively eluted in 300 mM NaCl, while shaking overnight at 4 °C. RNA was precipitated by addition of three volumes of ice-cold ethanol and recovered by centrifugation. The RNA pellet was resuspended in 200 µL nuclease-free water.

### Inter-strand ligation assay

Labeled RNA oligonucleotide ‘20.25’ was annealed to non-labeled complementary RNA oligonucleotide ‘19.1’ (5’-CGU ACG CGG AAU ACU UCG A-3’, Dharmacon, Thermo Scientific) in 30 mM HEPES KOH pH 7,5, 2 mM MgCl_2_ and 100 mM KCl and were heated to 95 °C for 2 min and subsequently incubated at 37 °C for 1 h. To test for inter-strand ligation, (3/5)th volumes of a reaction mixture (67 mM KCl, 2 mM MgCl_2_, 8,3 mM DTT, 5 mM ATP, 0,3 mM GTP, 53 units/mL RNasin RNase inhibitor, 43% glycerol) containing 17 nM radiolabeled RNA duplex were mixed with (2/5)th volumes of cell extracts (protein concentration 5 mg/mL) and incubated at 30 °C for 30 min. The reaction was stopped with equal volumes of 2x FA buffer and separated on a 15% denaturing urea polyacrylamide gel. Inter-strand ligation products were monitored by phosphorimaging, and band intensities was measured using ImageQuant (GE Healthcare) software and corrected by subtraction of background values.

### Pre-tRNA splicing assay

A PCR was performed using *S. cerevisiae* genomic DNA as template, a 5’ primer including the T7 polymerase promoter (5’-AAT TTA ATA CGA CTC ACT ATA GGG GAT TTA GCT CAG TTG GG-3’), and a 3’ primer (5’-TGG TGG GAA TTC TGT GGA TCG AAC-3’). The PCR product was sequenced and identified as yeast tRNA3-Phe^GAA^ (chromosome 13). The PCR product served as template for *in vitro* transcription using the T7 MEGAshortscript kit (Invitrogen, #AM1354) including 1,5 MBq [α-^32^P]guanosine-5’-triphosphate ([α-^32^P]-GTP, 111 TBq/mmol, Hartmann Analytic) per reaction. The body-labeled pre-tRNA was resolved on a 10% denaturing urea polyacrylamide gel, visualized by autoradiography and passively eluted from gel slices overnight in 300 mM NaCl. RNA was precipitated by addition of three volumes of ice-cold ethanol and dissolved to 0,1 μM in buffer containing 30 mM HEPES-KOH pH 7,3, 2 mM MgCl_2_, 100 mM KCl. To assess pre-tRNA splicing, one volume of 0,1 μM body-labeled *S. cerevisiae* pre-tRNA^Phe^ was pre-heated at 95 °C for 60 sec and incubated for 20 min at room temperature, and subsequently mixed with four volumes of reaction buffer (100 mM KCl, 5,75 mM MgCl_2_, 2,5 mM DTT, 5 mM ATP, 6,1 mM Spermidine-HCl pH 8,0 (Sigma Aldrich, # S4139), 100 units/mL RNasin RNase inhibitor (Promega, #N2515)). Equal volumes of this reaction mixture and cell extracts were mixed and incubated at 30 °C. At given time points, 5 μL of the mix were deproteinized with proteinase K (Invitrogen, #25530-049), followed by phenol/chloroform extraction and ethanol precipitation. Reaction products were separated on a 10% urea denaturing polyacrylamide gel, and mature tRNA and tRNA exon formation was monitored by phosphorimaging. Quantification of band intensities was performed using ImageQuant software (GE Healthcare).

### Kinetic trapping using FLAG-TRX overexpression cell lines

Trapping of TRX targets using FLAG-TRX overexpression construct was done as described before (Nakao et al., 2015). HeLa RIEP shTRX cells constitutively overexpressing FLAG-TRX CxxS were seeded into 15 cm^2^ plates and silencing of endogenous TRX by shRNA was induced with 2 µg/mL Dox for 3 days. Cells were left untreated, stressed for 3 min with H_2_O_2_, 15 min with menadione, for 4 or 8 h with 1,5 µg/ml Tm or 300 nM Tm. Cells were washed once and scraped in PBS supplemented with 20 mM NEM. After incubation of the cells in PBS with 20 mM NEM for 7 min, cells were pelleted for 5’ at 1000 rpm, and washed with PBS. The cell pellets were lysed in buffer 1 (25 mM TRIS, pH 7,4, 5 mM MgSO_4_, 150 mM NaCl, 0,3% Triton X-100) supplemented with cOmplete™ EDTA-free Protease Inhibitor (Roche, #05056489001) and 20 mM NEM, and incubated for 30 min under slight rotation at 4 °C. The lysates were cleared by centrifugation at 14 800 rpm for 10 min, 4 °C, protein concentrations were measured by Bradford assay and adjusted to 1-3 mg/mL in 800 µL total volume. 50 µL M2 FLAG-beads (Sigma Aldrich, #A2220-5ML) per 15 cm^2^ plate were washed three times in buffer 1 with 20 mM NEM, and added to the diluted cell lysate. Pulldown of FLAG-TRX variants with trapped proteins was performed for 3 h at 4 °C while rotating. Subsequently, beads were washed four times in wash buffer with increasing salt concentrations (buffer 1 with increasing NaCl concentrations: 150 mM NaCl, 250 mM NaCl, 400 mM NaCl or 500 mM NaCl) and four times in reverse order. Bound complexes were eluted in 50 µL 1 mg/mL FLAG-peptide dissolved in buffer 1 for 30 min, 4 °C, while shaking.

### Statistical analysis

To assess statistical significance in inter-strand ligation assay quantifications, mRNA expression levels (RT-qPCR) or FACS analysis we used unpaired or paired Student’s t test assuming unequal variances, with ∗, p < 0.05; ∗∗, p < 0.01; ∗∗∗, p < 0.005 using GraphPad Prism software.

### Calculation of IC_50_ values

For calculation of IC_50_ values quantification data from inter-strand ligation assays were normalized to the untreated sample of shCtrl cells. IC_50_ values were calculated by non-linear regression and sigmoidal concentration-response curve fit using GraphPad Prism software.

## Acknowledgements

We thank Igor Asanovic, Stefan Weitzer, Elif Karagöz and Tomas Aragon for help and advice. Flow cytometry was performed at Max Perutz Labs BioOptics FACS Facility.

## Funding information

Work at the Martinez lab was funded by the Medical University of Vienna, the “Fonds zur Förderung der wissenschaftlichen Forschung (FWF)” as Stand-Alone Projects (P34895 and P32011), the RNA Biology Doctoral Program and the SFB RNA Deco Program.

## Author contribution

D.J. designed and carried out most of the experiments. J.I. carried out FACS-based XBP1-splicing experiments and contributed to cell culture experiments. P.K. contributed to cell culture experiments. J.M. designed the experiments. D.J. and J.M. wrote the manuscript.

## Competing interests

No competing interests to be reported.

---

**Suppl. Figure 1, related to Figure 1:**
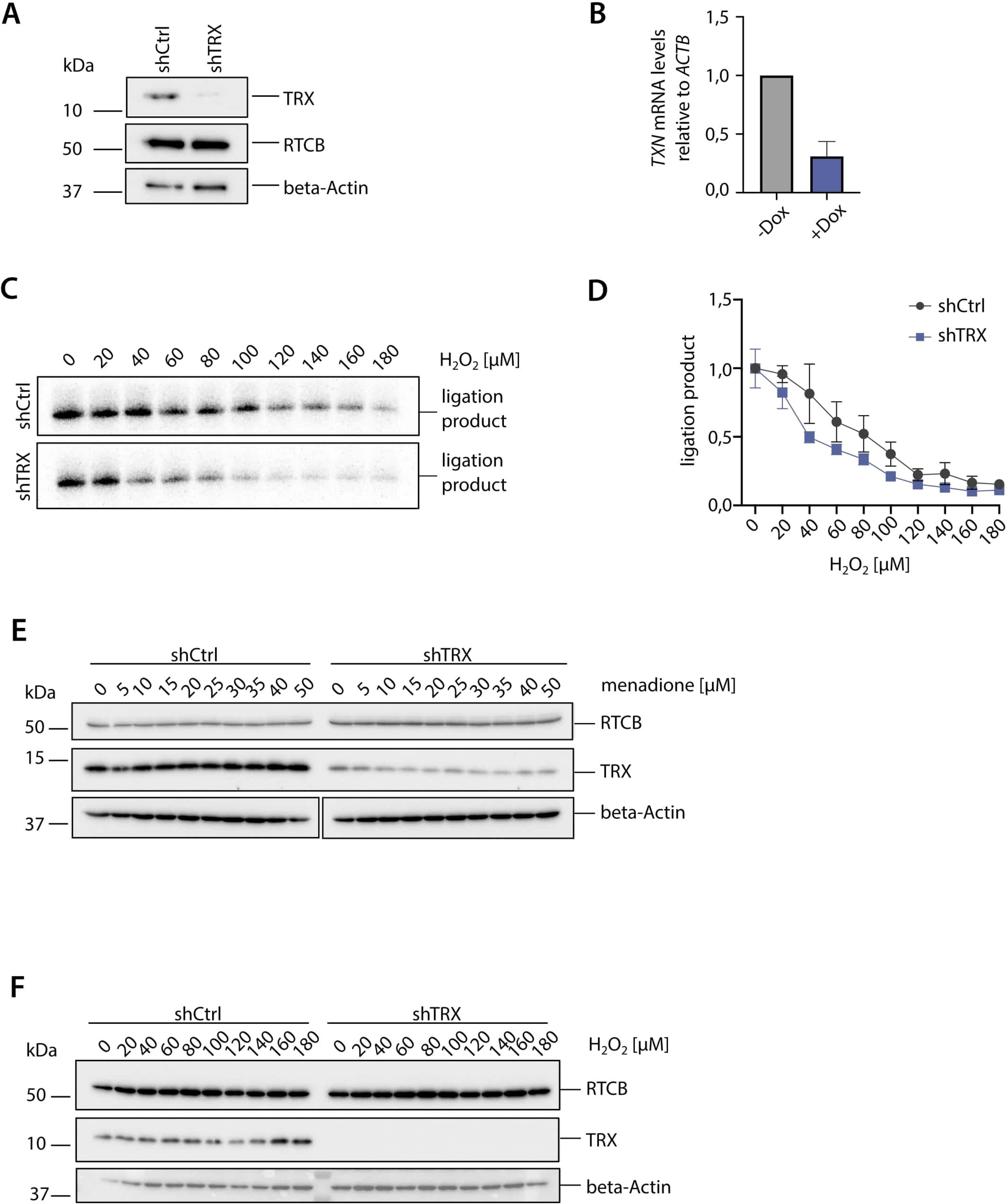
Depletion of TRX by shRNAs sensitizes the tRNA-LC to oxidative stress. A) HeLa cells expressing shTRX or shCtrl upon addition of 2 µg/mL Dox for 3 days were lysed and extracts were analyzed by western blotting using an anti-TRX antibody and anti-RTCB antibody. Levels of beta-Actin were assessed as a loading control. B) RNA was isolated from HeLa cells expressing shTRX, either induced with 2 µg/mL Dox or left uninduced for 3 days. Silencing of *TXN* mRNA levels relative to *ACTB* mRNA levels was assessed by RT-qPCR. C) ShCtrl and shTRX cells were treated with increasing concentrations of H_2_O_2_ for 1 h before harvest and lysis. Cell lysates were assayed for RNA ligation activity as in Fig. 1A. D) Quantification of band intensities from Suppl. Fig. 1C. The substrate conversion rate was calculated as in Fig. 1B, with n=3. E) Levels of RTCB and TRX were assessed in shTRX and shCtrl cell lysates after treatment with increasing concentrations of menadione for 1 h by western blotting using anti-RTCB and anti-TRX antibodies. Levels of beta-Actin were used as a loading control. F) Levels of RTCB and TRX were assessed in shTRX and shCtrl cell lysates after treatment with increasing concentrations of H_2_O_2_ for 1 h as in Suppl. Fig. 1E.

**Suppl. Figure 2, related to Figure 2:**
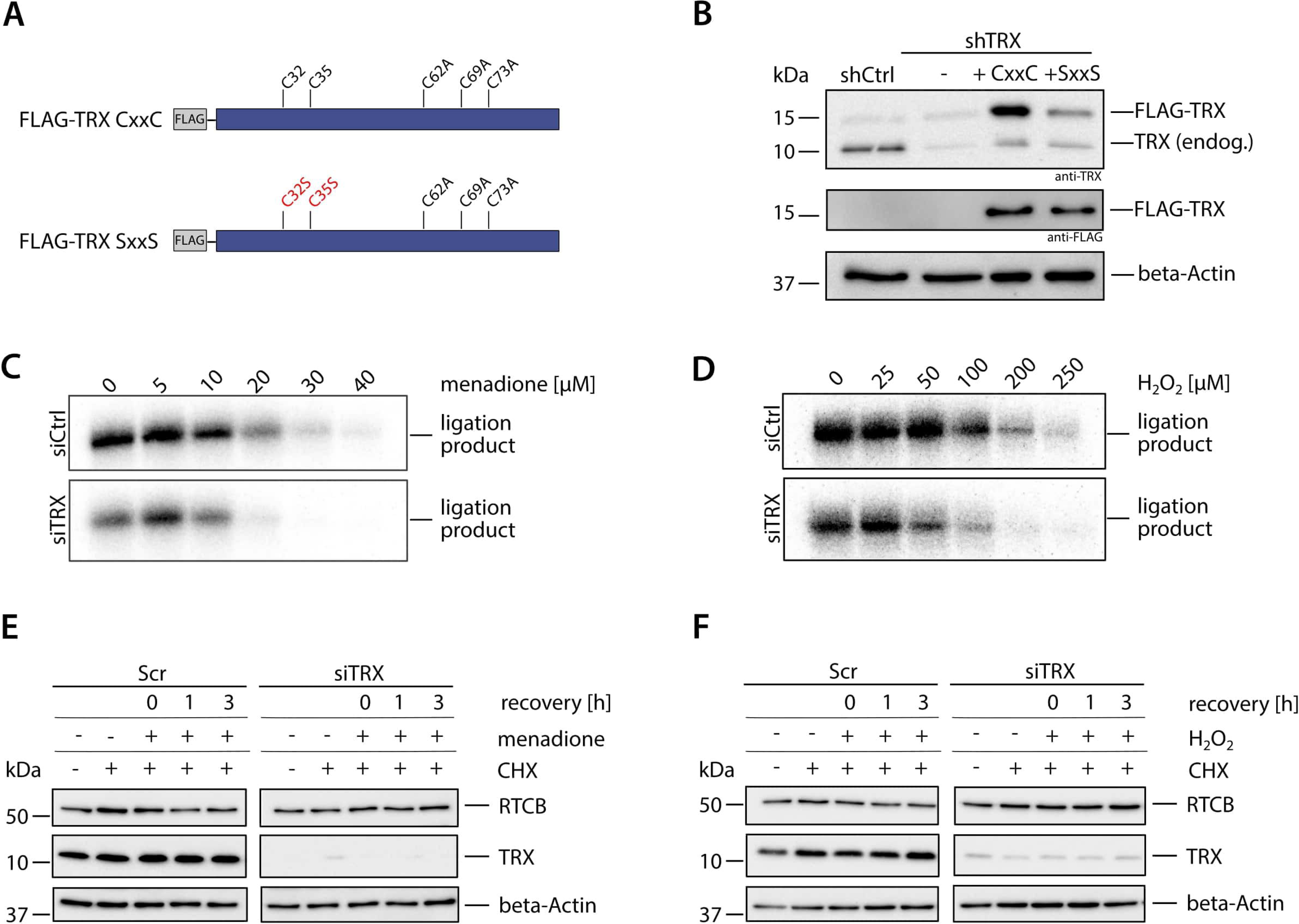
Catalytically active TRX is required to maintain tRNA-LC activity during oxidative stress and for its re-activation after oxidative stress levels decrease. A) FLAG-tagged variants of TRX CxxC and SxxS used for overexpression and testing of tRNA-LC activity after menadione treatment. Mutations of the active site in the variants are depicted in red. Three conserved cysteines close to the C-terminus were mutated to alanine to avoid involvement of the cysteines in redox reactions (Watson et al. 2003; Schwertassek et al. 2007). B) Western blot analysis of endogenous TRX in shCtrl, shTRX and rescue cell lines thereof overexpressing FLAG-TRX CxxC or SxxS double mutant. FLAG-tagged TRX CxxC or SxxS double mutant were only expressed in shTRX cells. Anti-TRX or anti-FLAG antibodies were used, and beta-actin levels were assessed as a loading control. Note: the anti-TRX antibody does not recognize FLAG-TRX SxxS double mutant as efficiently as FLAG-TRX CxxC. C) HeLa cells were transfected with siRNA pools targeting *TXN* mRNA (siTRX) or a control siRNA (siCtrl) for 3 days. Cells were treated with increasing concentrations of menadione for 1 h and RNA ligase activity was assessed as in Fig. 2A. D) HeLa cells were transfected with siRNA pools targeting *TXN* mRNA (siTRX) or a control siRNA (siCtrl) for 3 days. Cells were treated with increasing concentrations of H_2_O_2_ for 1 h and RNA ligase activity was assessed as in Fig. 2A. E) RTCB and TRX protein levels were assessed by western blotting in cell lysates of siCtrl and siTRX transfected cells after treatment with 30 µM menadione and different recovery times in CHX-containing medium using anti-RTCB and anti-TRX antibodies. Beta-actin levels were assessed as a loading control. F) RTCB and TRX protein levels were assessed by western blotting in cell lysates of siCtrl and siTRX transfected cells after treatment with 125 µM H_2_O_2_ and different recovery times in CHX-containing medium as described in Suppl. Fig. 2E.

**Suppl. Figure 3, related to Figure 4:**
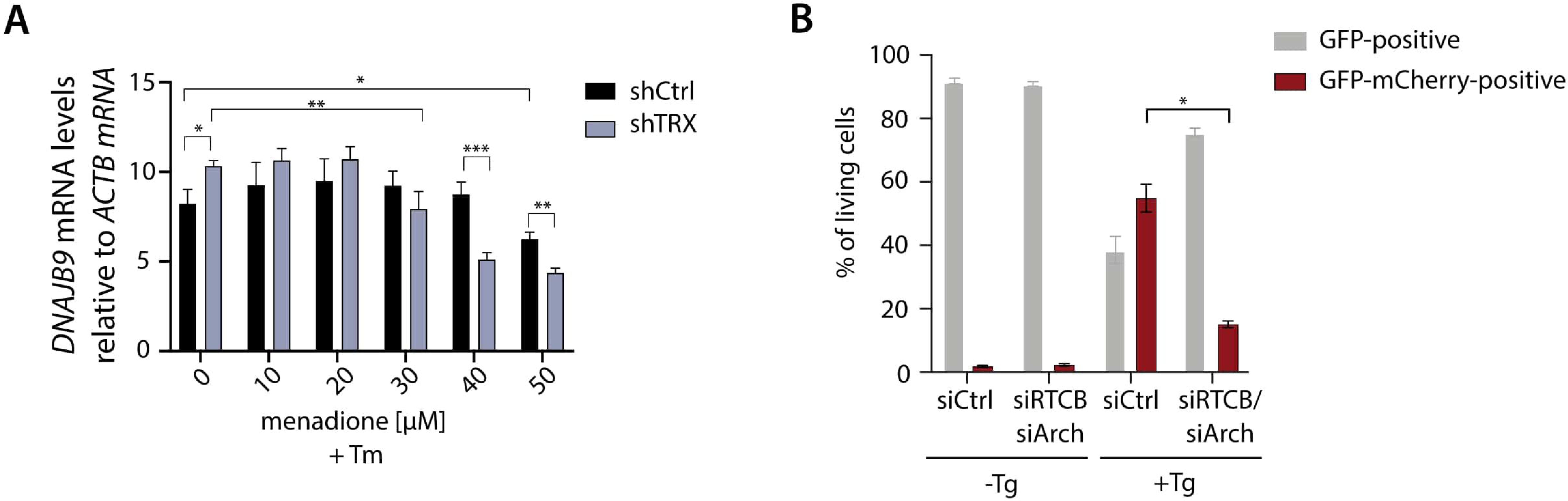
Depletion of TRX impairs physiological functions of the tRNA-LC. A) ShCtrl and shTRX cells were treated with indicated menadione concentrations and 1,5 µg/mL Tm for 4 h before harvest. RT-qPCR analysis of *DNAJB9* levels was performed as described in Fig. 4C, with n=9. B) HEK293 FITR cells expressing GFP-XBP1^intron^-mCherry were transfected with siRNA pools targeting *RTCB* and *Archease* mRNA (siRTCB/siArch) or control siRNA (siCtrl) for 3 days. Expression of the reporter construct was induced for 5 h using 2 µg/mL Dox and UPR was induced with 300 nM Tg. Splicing of the intron (generation of GFP-mCherry fusion protein) was assessed as in Fig. 4F, with n=3.

## Notes

### Competing Interest Statement

The authors have declared no competing interest.

